# Spectrast: a multiscale spatial statistics toolkit for single-cell spatial transcriptomics

**DOI:** 10.64898/2026.11.13.766464

**Authors:** Ian Gingerich, H. Robert Frost

## Abstract

Spatial transcriptomics resolves gene expression in tissue context, but the neighborhood statistics used to quantify spatial organization scale poorly to imaging-based datasets with hundreds-of-thousands of cells and thousands of genes. We recast the standard spatial-autocorrelation statistics as convolutions on a rasterized expression lattice and evaluate them with the fast Fourier transform, giving a single engine whose cost is governed by the neighborhood radius rather than the cell count. The engine computes each statistic across a continuous range of spatial scales from one shared transform, extends naturally to a differential test of spatial reorganization between tissue states, and runs unchanged on multicore CPUs or GPUs.

## 1 Introduction

Spatial statistics are used to describe a phenomenon, such as a signal, variable occurrence, or pattern, that is spatially distributed in some real-world coordinate space. In the 1950s and 1960s, these ideas were formalized quantitatively through mathematical definitions proposed by the likes of Moran [1] and Geary [2], and later work introduced statistics for clustering of high values [3] and local measures that describe each location rather than summarize the entire field study area [4, 5]. Around the same time, point-process methods studied the arrangement of events, rather than values measured at fixed locations [6–8]. These approaches now form complementary parts of a mature spatial statistics field of study [9, 10]. These statistics work by defining a spatial weights matrix whose elements capture a function of the distance between profiled locations, e.g, inverse-squared Euclidean distance. This weights matrix is usually thresholded at a specific radius for computational performance and memory considerations. The radius used as the threshold, often a fixed default in many analytic tools, can be thought of as a hypothesis about the scale at which the spatial phenomena operate. Literature in ecology and geospatial sciences has shown that these measures can be sensitive to radius selection, and the proposal of a multiscale spatial statistic has been presented as an alternative to a single-radius measure [11, 12]. This concept, called a scalogram by analogy with the scale-resolved energy of a wavelet transform [13], shows the change in a spatial statistic as a function of increasing radius. Scalograms can uncover spatial patterns that would otherwise be missed at a fixed radius, e.g., a feature organized into small, repeated foci and one organized into a single broad territory can return the same value at one radius but separate cleanly in a scalogram [11].

With the rise of spatial profiling technologies such as spatial transcriptomics (ST) [14–16], spatial analysis has also become commonplace in biology. Identifying spatially variable genes is a common step in the ST analytic pipeline, and various methods commonly use spatial statistics through the formulation of a spatial weights matrix computed of cell coordinates [17–25]. Spatially variable gene detection, spatially weighted pathway scoring [26], and cell–cell interaction inference [27–33] are common. However, unlike datasets in the ecology or geospatial fields, ST datasets are inherently high-dimensional. Instead of computing a scalogram on a few variables, ST datasets measure thousands of genes, and, in some technologies, the whole transcriptome (e.g., the 19k human gene probes used in the 10x Atera platform), on increasingly large tissue sections of over 1 million cells. Constructing a spatial statistic scalogram on high-dimensional ST data is computationally intractable with current tooling, and as such, a single radius or *k*-nearest-neighbor graph is used when constructing the spatial weights matrix for these methods.

Existing methods account for scale in different, often narrowly defined, ways. SPARK-X computes multiple coordinate transforms with scale parameters based on coordinate quantiles, but the results are combined into a single p-value rather than reported as a scalogram [19]. CRAWDAD reports multiscale relationships restricted to cell type labels, not genes [34], and Kontextual highlights how spatial relationships depend on the frame of reference used to measure them [35]. Other tools report multiscale geometric and spatial statistics [36] or identify transcript clusters at subcellular, cellular, and tissue scales [37]. These approaches all address scale-dependent questions but do not provide a full scale profile or scalogram for every gene in the dataset.

To address these computational limitations, we propose Spectrast, which solves this tractability problem in two ways. First, we turn irregular ST data into a regular lattice through rasterization. Rasterization of ST is not new, but has been only been employed to speed up the canonical pre-processing pipeline [38]. We improve upon a simple rasterization strategy by checking the grid approximation and refining cell-level estimates where needed in a rasterization-fusion approach. Second, once the ST data is rasterized, we can take advantage of convolution with an appropriate kernel, which can be performed efficiently using a two-dimensional Fast Fourier Transform (FFT). This enables the efficient computation of multiscale spatial statistics on ST data as well as inference on multiscale gene organizational changes between conditions. Spectrast enables several spatial measures in a single toolkit, making it practical to quantify gene organization from local neighborhoods to larger tissue regions. Its efficiency advantage is greatest at large radii distances, where graph-based calculations become computationally intractable for large ST datasets. The resulting scalograms reveal patterns that a single radius measurement would miss. We summarize each gene’s curve by its peak amplitude and characteristic scale, and develop a permutation-based test to identify changes in spatial organization between conditions. Together, Spectrast makes scale an explicit part of the ST analysis, allowing researchers to ask whether a gene is spatially organized, over what distances this organization occurs and how that organization changes across conditions.

## 2 Methods

The following sections summarize Spectrast’s computational approach, spatial statistics, and statistical tests supported. We first describe spatial statistics, how they can be computed efficiently across neighborhood sizes via convolutions, then explain grid construction, additional spatial measures, and inference. Full definitions, implementation details, parameter defaults, and validation procedures can be found in the Supplementary Methods.

### 2.1 Spatial statistics as quadratic forms

Spatial statistics describe how measured values, such as gene expression, are arranged or organized across locations in geographic space. Global Moran’s *I* summarizes whether nearby values are similar, Geary’s *C* measures differences between neighboring values, and Getis–Ord general *G* measures the spatial concentration of high values. These statistics use a spatial weights matrix *W* ∈ℝ^*n×n*^ in their computation, where *n* is the number of locations measured. Each entry *w*_*i,j*_ specifies how much location *j* contributes to the neighborhood summary at location *i*. Weights may be graded so that nearby locations have more influence than distant ones, or binary, so every location in the neighborhood contributes uniformly. A zero weight indicates no contribution and diagonal entries are set to zero to exclude self-contributions.

A hard neighborhood boundary is not required when formulating *W*, but weights are commonly restricted to nearby locations for computational efficiency, using a distance threshold or a fixed number of nearest neighbors. Spectrast uses a radius-based neighborhood as a hard cutoff. Distinct locations within a radius *r* are neighbors and contribute equally, whereas those outside the radius contribute zero. Depending on the statistic, these connections are either used directly or rescaled. We write *W*_*r*_ for this radius-dependent weights matrix; statistic-specific normalizations are described in Supplementary Section S2.

While these spatial statistics give different insights into the data, the way they are computed is very similar, with differences only in how the spatial weights are formed, which values are used in comparison, and how the result is scaled so it can be interpreted. These similarities are captured in a shared quadratic form equation:

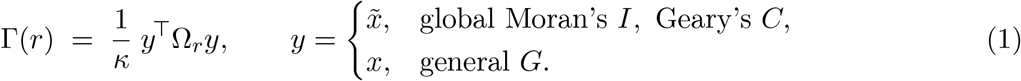

Here, *x* ∈ ℝ^*n*^ contains the measured values at *n* locations, such as one gene’s expression across cells. The vector 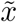 contains the standardized values, with zero mean and unit variance, so that 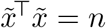. The operator Ω_*r*_ is the statistic-specific form of the spatial weights *W*_*r*_ and *κ*(*r*) is the statistic-specific normalization [39] (Fig. 1a). Structured in this way, applying Ω_*r*_ to the gene expression values becomes the only step that involves distance/location information. That step can be carried out as a convolution which is described in section 2.2. The exact per statistic operator construction (Ω_*r*_) and normalizations (*k*(*r*)) are given in Supplementary Section S2 and Supplementary Fig. S1.

**Figure 1:**
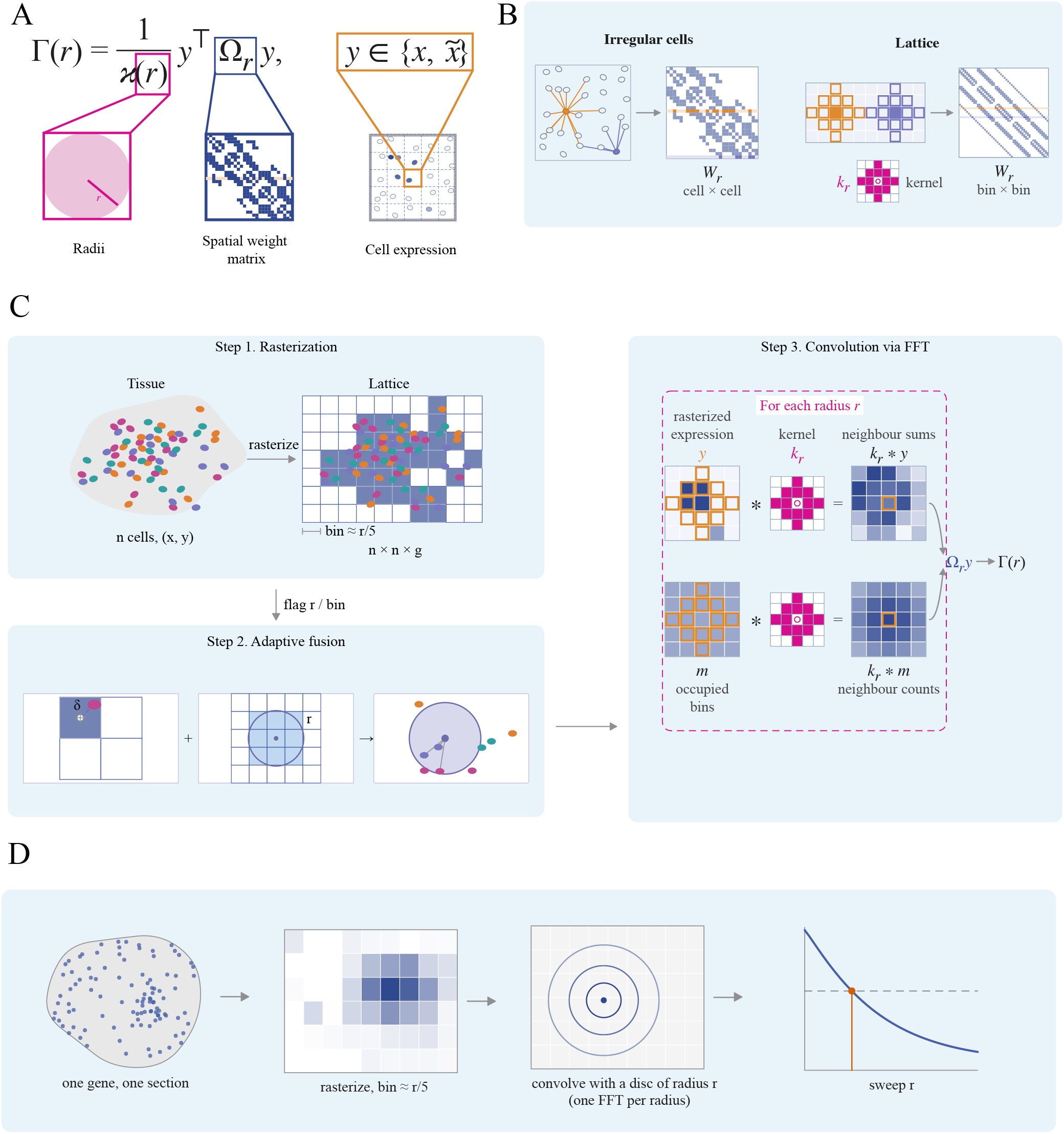
Algorithm overview. **A.** The general quadratic form equation for global Moran’s I, Geary’s C, and General G. **B**. Schematic illustrating the direct formulation of the spatial weights matrix from cell positions vs the shared spatial kernel available for rasterized data. **C**. Overview of the Spectrast’s key steps; rasterization, correction of distortion, and convolution via FFT. Both routes are drawn at the same radius *r* around the same example cell and bin (orange). **D**. Schematic illustrating how a single spatial statistic turns into a scalogram.

### 2.2 Fast multiscale computation and spatial summaries

On an irregular set of points, such as those of imaging based ST data, every location has its own set of neighbors, so each row of *W*_*r*_ is unique and must be applied individually. However, on a regular grid, or lattice, locations are evenly spaced, and the neighborhood is the same everywhere, meaning that *W*_*r*_ holds the same pattern of weights, with only the center being shifted. We handle edge effects by excluding bins with zero neighbors, after zero padding. This repeated pattern is explicitly constructed into an operator called a kernel by Spectrast. The kernel is a small array for which entries are the weight each surrounding location contributes to a specific location’s neighborhood sum. By convolving this kernel across the regular grid, this spatial weighting can be applied to every location without explicitly building *W*_*r*_ (Fig. 1b). The convolution theorem allows these sums to be calculated by multiplying the 2D discrete Fast Fourier transforms (FFT) of the data and kernel, then transforming the resulting product back to regular space. Spectrast uses the FFT for this calculation, avoiding the need to construct and multiply a large spatial weights matrix. Notably, the data’s forward transform can be reused across neighborhood radii, with only the kernel changing (Fig. 1c, right side) allowing evaluation of spatial statistics across many neighborhood sizes rather than being limited to a single radius.

The values of a given statistic evaluated across multiple radii result in a spatial statistic curve. Following the nomenclature of signal processing, ecology and geospatial science, we call this curve a *scalogram* [12, 13], which shows how spatial organization of a gene changes from local to broader tissue scales. We summarize it by its peak height, the largest statistic value across the tested radii, and its characteristic scale, the radius where the curve crosses halfway between its maximum and minimum after applying a common orientation across genes. For global Moran’s *I*, smaller characteristic scales indicate finer spatial organization and larger scales indicate broader spatial organization.

Scale estimates depend on the tested radius range. We withhold estimates from weak curves or curves without a suitable midpoint crossing, and flag estimates that may reflect the chosen radii rather than tissue structure. The next section describes how ST measurements are placed onto a grid. Convolution, scale estimation, and quality-control details are given in Supplementary Sections S2.3, S3.3–S3.4, and S4, and Supplementary Fig. S2.

### 2.3 Rasterization of ST data

To utilize convolution-based computations, Spectrast places irregular imaging based ST measurements on a regular grid, averaging values for cells assigned to the same bin and recording which bins contain cells. When forcing ST data to a grid, the selected bin width has a direct relationship with the smallest radii a scalogram can use, as radii smaller than a bin width would not be resolvable. To balance these two values, we select the smallest neighborhood radii for a scalogram from the pooled median cell spacing across the tissue, then choose a grid resolution fine enough to resolve that radii. However, rasterization introduces some amount of spatial distortion, so we compare cell-level statistics on the selected grid with those on a grid twice as fine. Values that vary beyond a set limit are recomputed directly from the spatial coordinates, using each flagged cells true neighbors within radius *r*. This combines fast convolution computation with direct spatial graph refinement only where needed (Fig. 1c, bottom right). Grid construction, parameter defaults, and refinement checks are described in Supplementary Section S4 and Supplementary Figs. S3 and S4.

### 2.4 Additional measures of spatial organization

In addition to the classic global Moran’s I, Geary’s C and Getis-Ord G statistics, Spectrast computes three additional spatial measures: local spatial heteroscedasticity (LOSH), focality, and spectral texture. LOSH summarizes local variance or dispersion, focality measures whether positive spatial autocorrelation is broadly distributed or concentrated in a few regions, and spectral texture uses the Fourier power spectrum to distinguish fine-grained patterns from smoother, broader patterns. These measures are evaluated on the same length-scale axis as the classical statistics, although spectral texture uses frequency bands rather than spatial neighborhoods. Definitions and calculations are given in Supplementary Section S2.4 and Supplementary Fig. S1.

### 2.5 Statistical inference

Spectrast tests whether genes show statistically significant spatial organization and whether that organization differs between conditions. For individual genes, it supports three approaches: analytic, surrogate, and permutation nulls. The analytic method uses known null means and variances for Moran’s I, Geary’s C, and general G, and then assumes a Gaussian distribution to estimate p-values. This is fast but can be inaccurate in the tails. Surrogate methods estimate null distributions from a subset of permuted genes, providing a faster approximation for gene ranking. Permutation tests shuffle values across cells and recompute the statistic, providing calibrated p-values for all six statistics. Shuffling can also be restricted within groups defined by cell density or another covariate to test for additional associations. Supplementary Table S3 lists the methods available for each statistic.

To compare statistics between conditions, we test differences in scalograms [40]. When users have multiple biological replicates, Spectrast shuffles condition labels across independent experiment units or swaps labels within paired units. Repeated samples from the same unit are averaged before testing. In cases with only one sample per condition, we instead create cell partitions within each sample to assess the difference. These partitions do not replace biological replication, and the test has limited power and its conclusions apply only to the measured samples, not to differences across subjects. The differential test accounts for searching across radii and corrects for multiple testing across genes using the Benjamini–Hochberg procedure. The gene level test indicates whether conditions differ somewhere along the scalogram, but to show where along the curve the difference occurs we construct simultaneous null bands around the expected zero-difference curve [41]. These bands describe the range of differences expected under the permutation null across the whole scalogram rather than confidence intervals around the observed values [8]. Radii where the difference curve leaves this band helps identify the scales that contribute to the unadjusted gene-level test. Spectrast also reports the smallest possible p-value given the sample size. Null models, experimental designs, and validation are described in Supplementary Sections S5–S6 and Supplementary Figs. S5 and S6.

### 2.6 Datasets

We used publicly available and in-house spatial transcriptomics datasets spanning five platforms, 10x Visium HD, 10x Xenium, Vizgen MERFISH, Stereo-seq, and 10x Atera whole-transcriptome imaging, and gene-panel sizes from 300 to ~ 18,000 genes. Eight datasets were used for benchmarking runtime, memory, and accuracy against reference implementations; the differential and descriptive biological analyses use the Trem2/5xFAD MERFISH, Lymph node Xenium datasets. Dataset accessions, post-quality-control cell and gene counts, and the complete quality-control, normalization, and dimensionality-reduction pipeline are given in Supplementary Section S1 (Table S1).

### 2.7 Benchmarking

We benchmarked Spectrast against graph and distance-based reference implementations across eight datasets spanning five ST technologies. Both approaches used the same radii, from 50 to 1600 *µ*m. We measured computation time and peak memory under the same single-threaded settings. Datasets, reference implementations, and measurement calculations are described in Supplementary Section S7.

### 2.8 Software

The toolkit is **SpectraST** 0.2.0 (this work), evaluated against **squidpy** 1.8.1 [23], with **scanpy** 1.11.5, **anndata** 0.12.10, **numpy** 2.4.4, **scipy** 1.17.1 [42], **scikit-learn** 1.8.0, and **numba** 0.65.0 under **Python** 3.11.

## 3 Results

### 3.1 Spectrast reproduces standard spatial statistics and accelerates large-neighborhood analysis

We benchmarked Spectrast across eight ST datasets that span five ST platforms (Table 1). Field standard spatial statistic implementations explicitly construct a spatial weights matrix and use it to calculate neighborhood sums. As the radius grows, more neighboring cell pairs sums need to be computed. Spectrast sidesteps this issue, computing these sums by FFT-based convolution on a grid, avoiding the weights matrix entirely and making the cost depend mainly on grid size rather than the number of neighboring pairs (Section 2.2; Fig. 1b). See SI S7 for benchmarking details.

**Table 1:** Benchmark datasets. Counts are reported after quality control. Acquisition, processing and accessions are given in Supplementary Section S1 (Table S1).

| Tissue | Platform | Cells | Genes |
|---|---|---|---|
| Mouse olfactory bulb | Stereo-seq | 19,109 | 14,376 |
| Mouse brain | 10x Xenium (5K) | 63,173 | 5,006 |
| Mouse brain | Vizgen MERFISH | 81,452 | 1,122 |
| Human breast cancer | 10x Atera (WTA) | 169,186 | 18,028 |
| Mouse brain | 10x Visium HD | 393,434 | 19,059 |
| Human ovarian cancer | 10x Visium HD | 439,300 | 5,000 |
| Human breast cancer | 10x Xenium (5K) | 565,916 | 541 |
| Human cervical cancer | 10x Atera (WTA) | 709,888 | 18,027 |

Spectrast closely matched reference calculations for all eight sections. Figure 2c shows the Atera Breast section accuracy plots for the top 10,000 most highly variable genes, and the remaining sections are shown in Supplementary Fig. S1. Across sections, median correlations were 0.99996 (95% CI 0.99989–0.99999) for global Moran’s *I*, 0.99995 (0.99986–0.99999) for Geary’s *C*, 0.99994(0.99973– 0.99998) for general *G*, and 0.99983 (0.99969–0.99998) for focality, with minima of 0.9990, 0.9985, 0.9985 and 0.9982 respectively. LOSH correlations were lower, with a median of 0.9683 (0.9191– 0.9860) and a range of 0.908 to 0.989 across sections. This was expected as this comparison is computed on the local cell rather than global level, comparing approximately 2.15 billion cell–gene values, including 1.69*×*10^9^ in the section shown. Global gene-level summaries such as global Moran’s I tend to average out local rasterization error, resulting in high correlation compared to the reference, where LOSH values highlight such errors.

**Figure 2:**
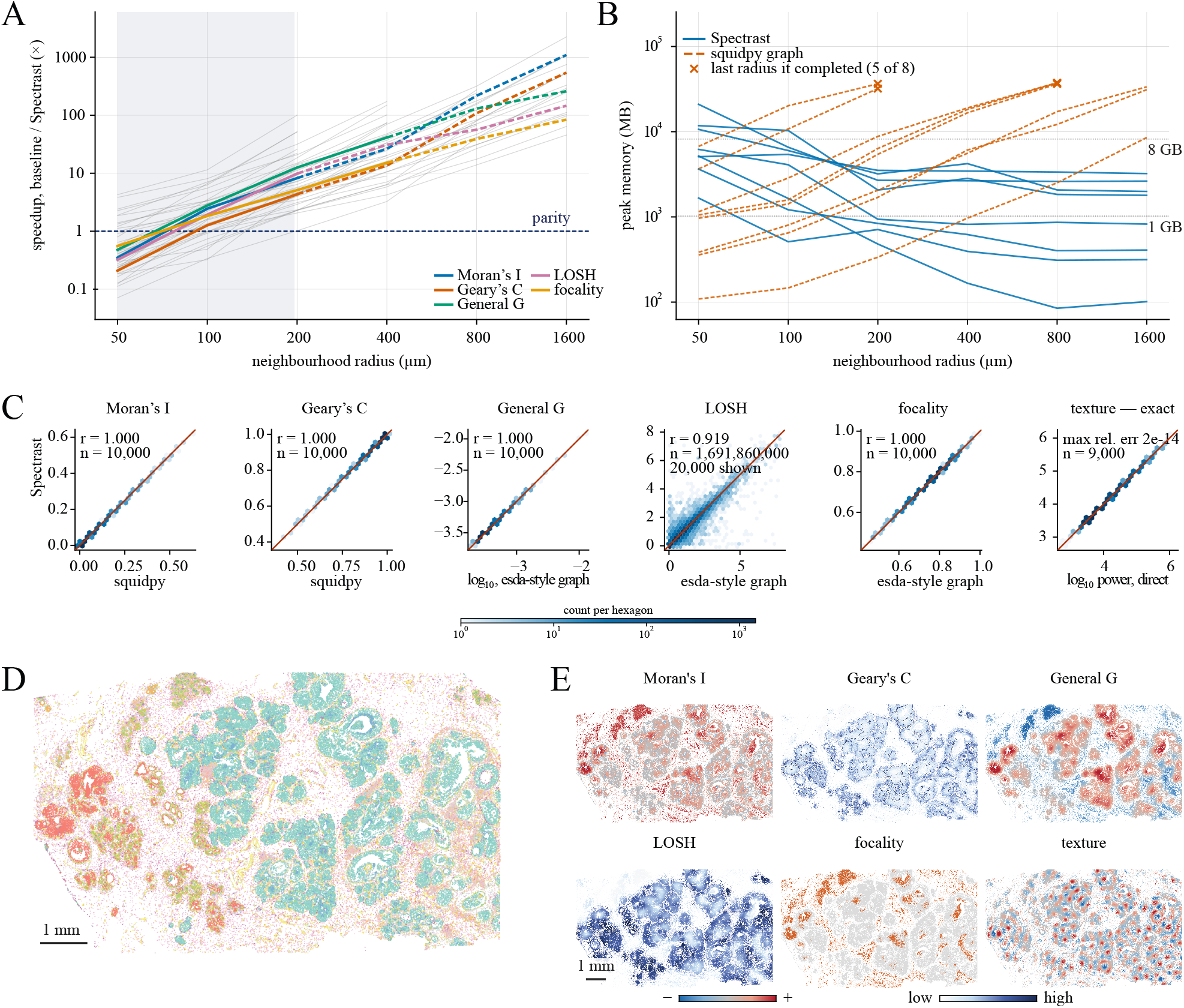
Spectrast reproduces standard spatial statistics and enables multi-scale analysis. **A.** Relative speedup of Spectrast over reference implementations for the five spatial statistics with a matched neighborhood baseline. Spectral texture is absent, as no neighborhood implementation exists to time it against. Gray lines represent the 40 statistic *×*dataset combinations over the eight datasets used for benchmarking, colored lines show the median values. The horizontal dashed line marks parity between the two implementations. **B**. Peak memory consumption of Spectrast (blue, solid) vs squidpy’s neighborhood graph (orange, dashed) for global Moran’s I, one line per dataset, over the same radius sweep. X’s mark the last radius the graph completed before OOM errors (48 GB). **C**. Spectrast and reference implementation spatial statistic values plotted against each other per gene (global Moran’s I, Geary’s C, General G, focality), per cell (LOSH), and per gene and band (spectral texture), on the Atera breast section. **D**. Atera breast dataset colored by cell type. **E**. Illustrative spatial scatter plots for the six spatial statistics visualized on the Atera breast tissue, drawn for NHERF1 at *r* = 223 *µ*m.

Spectrast’s speed advantage is dependent on neighborhood size, rather than being uniformly faster than reference implementations (Fig. 2a). At 50 *µ*m, all five statistics with graph-based timing references were slower on median, with speedups of only 0.21–0.55*×*. By 100 *µ*m every statistic was faster on median (1.26–2.77*×*), and the advantage compounded with radius, from a pooled median of 7.2 at 200 *µ*m to 276*×* at 1600 *µ*m. Spectrast was slower in 40 of the 196 radius–tissue–statistic conditions, all at 50 or 100 *µ*m. This crossover reflects the different computational costs of the two approaches. Graph-based calculations must represent and process more neighboring pairs as the radius grows. FFT-based convolution depends mainly on the grid size rather than the number of pairs within a neighborhood. Small neighborhoods may not justify an FFT approach, but larger neighborhoods increasingly favor convolution.

Memory usage also shows a radii-dependent difference between Spectrast and reference implementations (Fig. 2b). For global Moran’s *I* and Geary’s *C*, Spectrast’s median peak memory usage fell from 5.66 to 0.31 GB between 50 and 1600 *µ*m, while the reference increased from 0.99 to 30.7 GB.

The median reference-to-Spectrast memory ratio increased from 0.14 to 83 over that range. Across 192 memory-profiled conditions, the operating system terminated 18 reference runs on a 48 GB machine due to out-of-memory issues, while no Spectrast runs were terminated. All failures occurred at larger radii in the graph based implementation: four at 400 *µ*m, four at 800 *µ*m, and ten at 1600 *µ*m. Graph-based tools therefore remain competitive for small neighborhoods with Spectrast’s main advantage at broader tissue scales, for which the reference calculations become much slower and more memory-intensive, sometimes exceeding available memory. The crossover reflects how each method handles computing *W* at increased radius sizes. Graph-based implementations store each cell’s neighbors, whose numbers grow roughly with area as *πr*^2^. Larger radii require more compute and memory, approaching an *n × n* matrix amount of resources for very large neighborhoods. Spectrast by comparison does not store neighborhood pairs. Its kernel grows with radius, but the grid is fixed by bin width. For small neighborhoods, graphical representation is cheaper than running Spectrast’s transforms, and for larger neighborhoods, convolution becomes more efficient. Spectrast makes it practical to follow spatial patterns across scales rather than restrict analysis to a single small neighborhood.

### 3.2 Peak height and characteristic scale recover lymph-node organization

Spectrast’s memory and speed advantages make it practical to calculate a spatial statics scalogram across multiple radii showing how spatial organization changes with scale. Methods section 2.2, and SI 3.3 describes how these scalograms are decomposed into two values: characteristic scale and amplitude. We investigated whether the peak height and characteristic scale of a scalogram capture recognizable tissue structure. We analyzed a human lymph node (10x, Xenium) with 708,983 cells and a 4,624-gene panel, as this tissue contains clear anatomical compartments that provide an independent basis for comparison. The observed expression patterns were separated for these two measurements when plotted using coordinate scrambled controls (Fig. 3a). The scrambled genes are confined to short scales and small amplitudes, while measured genes extend an order of magnitude beyond in amplitude and twice as far along the scale axis. The position of each gene reflects spatial arrangement rather than expression distribution. Scale estimates were restricted to the top fifth of the genes by amplitude for visualization purposes: 923 genes spanning 26–178 *µ*m. Focality provided much weaker separation from the scrambled controls, so we therefore used it as a color scale to distinguish genes with similar peak heights and characteristic scales, rather than as a measure of signal strength by itself.

**Figure 3:**
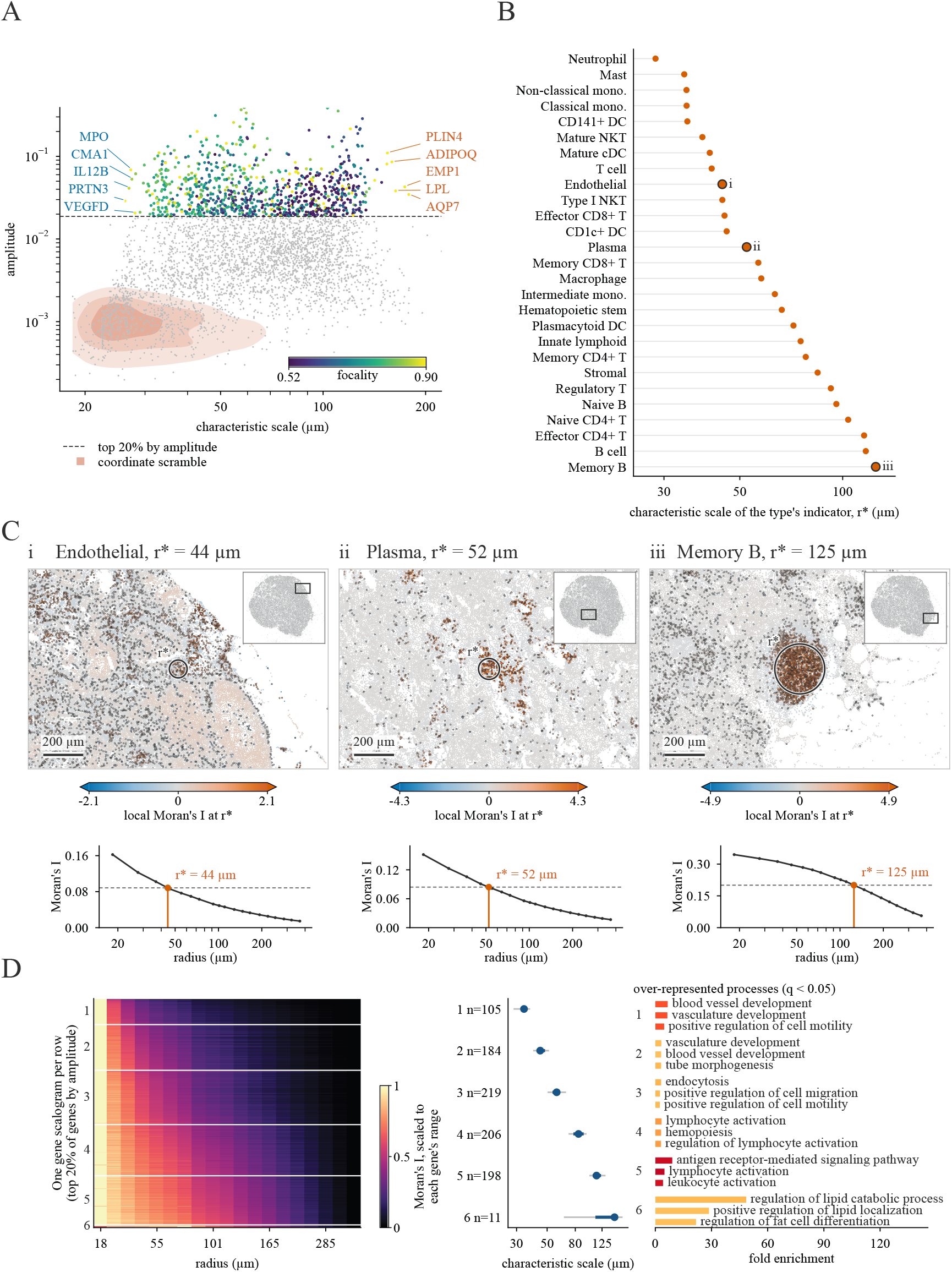
Peak height and characteristic scale recover the organization of a human lymph node. **A.** Scatter plot of the characteristic scale and amplitude for each gene’s global Moran’s I scalogram. Color scale indicates each focality values. Only genes in the top 20% of amplitude are colored (923 of 4,624). The tan area represents a kernel density estimate of the joint scale and amplitude null distribution, with each hue representing contours at 10%, 35% and 70% of the peak null density respectively. **B**. Characteristic scale for cell type indicator’s global Moran’s I scalogram, ordered from smallest to largest, for the 27 annotated cell types with at least 500 cells. Circled bubbles represent the cell types illustrated in panel C. **C**. Local Moran’s I spatial scatter plots for endothelial (left), plasma (middle) and memory B cells (right) computed at their characteristic scale radii, with circles matching these radii. Example scalograms for each cell type indicator shown below. **D**. Scalograms clustered by *k* = 6 k-Means clustering (left), scatter plot of cluster’s median characteristic scale (blue) and 5th–95th percentile range (gray) (middle), and barplot of Gene Ontology enriched terms for each cluster (right), showing up to three terms per cluster.

We applied the same analysis to cell type indicators, which allows cell types and genes to be compared on the same scale axis (Fig. 3b). Of all annotated cell types with at least 500 cells, 27 had supported scales that spanned 28–125 *µ*m. Neutrophils, mast cells, and monocytes had the smallest characteristic scales (28–35 *µ*m), followed by dendritic and T cells (35–46 *µ*m), plasma cells and macrophages (52–58 *µ*m), and naive, mature, and memory B cells (96–125 *µ*m). Note that Erythrocytes did not receive a scale estimate because their spatial autocorrelation was weak (591 cells; peak global Moran’s *I* = 0.004), and their scalogram curve had no midpoint crossing. Figure 3c shows the local spatial statistic at the characteristic scale of three cell types: endothelial cells (44 *µ*m), plasma cells (52 *µ*m), and memory B cells (125 *µ*m). Smaller scales capture tight or repeated structures, such as the fine endothelial vascular network, while larger scales capture broader regions, such as the wide memory B-cell follicle. We used k-means to cluster the curves of the top 20% of genes by amplitude into six groups, ordered by median characteristic scale. Each group was tested for over-representation of Gene Ontology (GO) biological processes using a one-sided hypergeometric test with BH FDR correction. All groups had significant GO term functional enrichment at FDR values *<* 0.05. These enrichment values were consistent with the expected lymph node spatial structure (Fig. 3d). Small-scale groups were enriched for vascular development and tube morphogenesis, matching the fine, branching structure of blood vessels. Broader-scale groups were enriched for lymphocyte activation and lipid metabolism, consistent with follicles and surrounding adipose tissue. The follicular gene group had a characteristic scale of 114 *µ*m, close to the independently measured B-cell scales of 96–125 *µ*m.

### 3.3 Spectrast detects spatial reorganization between conditions

Spectrast was also tested to see if it could detect differences in spatial organization between conditions by comparing the scalogram for each gene computed in both conditions (Fig. 4a). The test gives a p-value per gene as well as a null band that shows where along the radius sweep the conditions differ, if at all. See SI for details (Supplementary Fig. S5).

**Figure 4:**
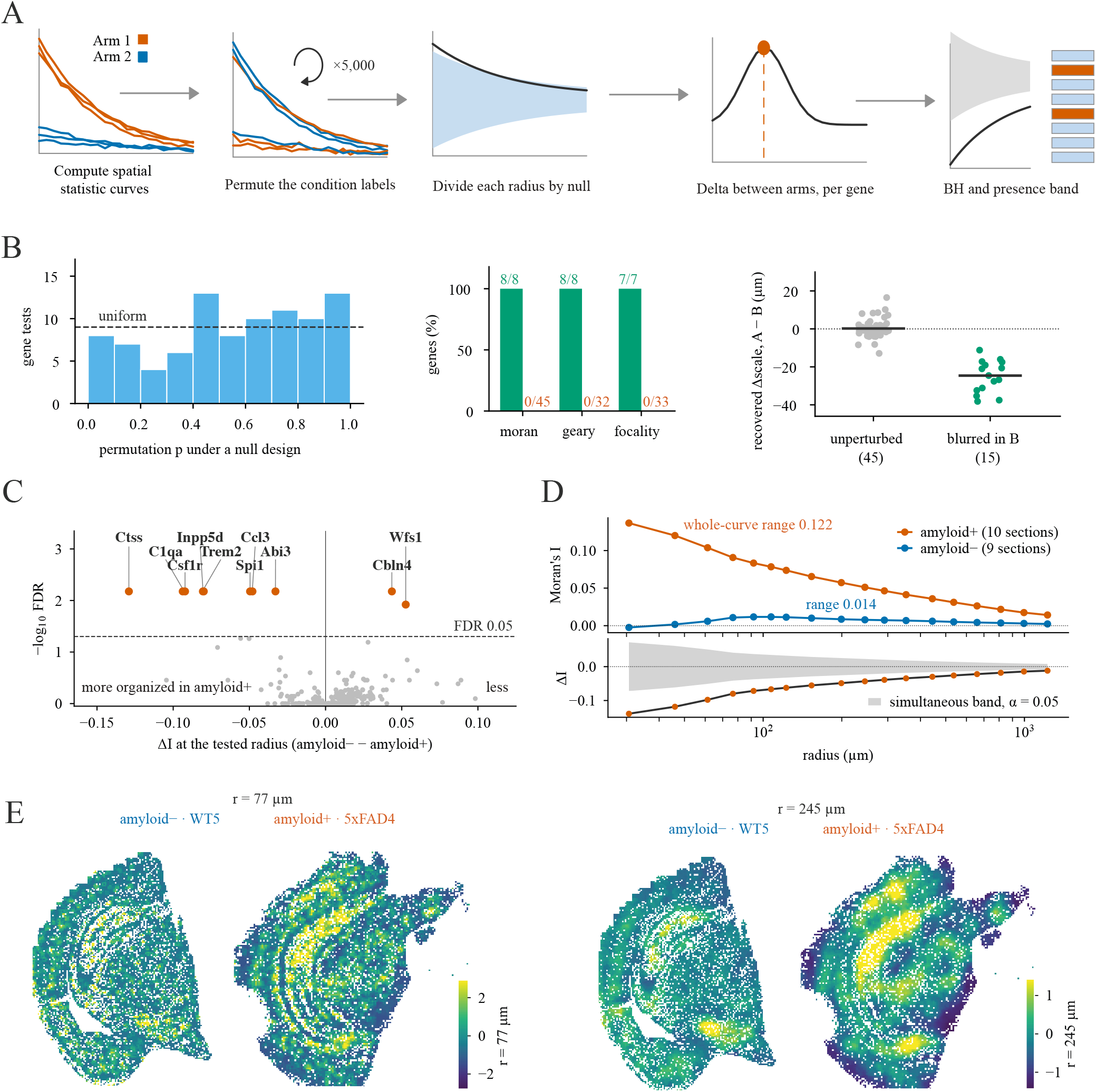
Spectrast detects spatial reorganization between conditions. **A.** Schematic illustrating Spectrast’s differential testing framework. **B**. Calibration on a Xenium 5K breast section, partitioned into arms (60 genes per experimental design). Bar chart showing p-values under artificially partitioned tissue sections (left; dashed line, the uniform expectation), bar chart depicting number of injected genes detected as differentially organized (green) vs false positive hits (orange) (middle), and paired scatter plot showing recovered scale shifts of artificially blurred genes (right). **C**. Volcano plot showing differentially organized genes (global Moran’s I) in the Trem2 MERFISH dataset: all 300 panel genes, of which ten reach FDR *<* 0.05 (orange). The horizontal axis is the difference in global Moran’s I at the tested radius, negative = more organized in amyloid-positive tissue. **D**. Example global Moran’s I scalograms for amyloid + Trem2 (orange), and amyloid - (blue) arms (top graph), and the delta curve and null band for this comparison (bottom graph). **E**. Example spatial scatter plots showing the neighborhood mean of the standardized Trem2 field for amyloid − and + arms smoothed at different radii.

#### 3.3.1 Calibration with perturbations

We evaluated false-positive control and sensitivity by creating artificial partitions of the human breast cancer (Xenium 5k) dataset. We tested the 60 most widely detected genes at five radii from 30 to 480 *µ*m on a 30 *µ*m grid, using 150 permutations. We used six different experimental designs to test that Spectrast did not generate false positive results, including balanced and unbalanced partitions, value scrambling, spatial smoothing, and scale recovery (Supplementary Section S6 and Supplementary Fig. S6).

Figure 4b summarizes false-positive control, scramble detection, and scale recovery. All significance is assessed on permutation *p*-values corrected across genes by the Benjamini–Hochberg procedure, at a threshold of FDR *<* 0.05. Across three partitions, an equal split, a 70*/*30 split, and a four-way split grouped into two pseudo-replicates per condition, no gene reached FDR *<* 0.05 on the global Moran axis. 3.3% of tested combinations had raw *p*-values *<* 0.05. A further 273 tests using Geary’s *C* and focality also produced no results at FDR *<* 0.05. For sensitivity tests, we perturbed 25% of genes in one arm. Value scrambling was detected at FDR *<* 0.05 for every eligible perturbed gene: 8 of 8 for global Moran, 8 of 8 for Geary, and 7 of 7 for focality, with none of the 110 accompanying unperturbed gene–test combinations flagged by Spectrast (Fig. 4b). Gaussian smoothing (*σ* = 120 *µ*m) produced the expected direction of scale change for all 15 perturbed genes (Fig. 4b). The median contrast was −24.6 *µ*m for perturbed genes and +0.2 *µ*m for unperturbed genes, with the negative sign reflecting the unsmoothed-minus-smoothed comparison.

#### 3.3.2 Spatial reorganization in the Trem2/5xFAD cohort

We applied Spectrast’s differential test to a *Trem2* ^R47H^/5xFAD mouse-brain MERFISH cohort, using replicate-label permutations and a radius sweep of 31–1226 *µ*m. Out of 300 panel genes, 10 showed differential spatial organization at FDR *<* 0.05 and 14 at FDR *<* 0.10 (Fig. 4c). These included eight genes associated with microglial biology and disease responses: Trem2, Ctss, Abi3, C1qa, Csf1r, Inpp5d, Ccl3, and Spi1, each with unadjusted p-values of 2 *×* 10^−4^ and FDR = 0.0067. In contrast, no gene showed differential mean expression at FDR *<* 0.05; the smallest expression FDR was 0.0514. Trem2 illustrates this change on treatment (Fig. 4d,e). Its scalogram was near zero across all radii in amyloid-negative tissue but reached 0.137 amplitude at the smallest radius in amyloid-positive tissue. The difference exceeded the null band at all 18 tested radii, indicating the emergence of spatial organization rather than simply a shift in an existing pattern. These changes can be seen by the spatial scatter plots in Fig. 4e.

## 4 Discussion

### 4.1 A shared framework for scale-resolved spatial statistics

Spectrast’s contribution is to make spatial statistics practical across a range of radii distances. Our rasterization and FFT-based approach reduces the repeated work needed for a radii sweep, allowing researchers to examine how spatial organization changes with distance rather than relying on a single distance to threshold the spatial weights matrix when computing these statistics. The importance of this choice has been reported in ecology and geostatistics [11, 12]. This computational advantage depends on the radius, and our benchmarks show that reference implementations outperform Spectrast below~ 100 *µ*m. At larger radii, Spectrast becomes increasingly advantageous, with a computational cost that is orders of magnitude below reference implementations at the largest tested radii. We therefore recommend Spectrast for multiscale sweeps, not as a faster replacement for existing tools in all cases. Spectrast achieves this acceleration while maintaining close agreement with reference calculations across spatial statistics and datasets tested. The biological examples highlight known lymph-node organization, and show genes that are reorganized between conditions in the differential test.

### 4.2 Future directions

Spectrast’s rasterization and convolution-based approach has a natural extension to bivariate spatial statistics that quantify the relationships between two spatially-distributed variables, e.g, genes or cell types. This includes cross-correlation statistics and functions, and co-localization measures. In the context of ST data, a natural application is cell-cell interaction analysis at the level of ligand-receptor or cell type pairs [27]. A scale-resolved cross-correlation analysis can be used to identify the distance at which a given ligand and receptor pair exhibit the strongest spatial association [31–33]. Similar analyses can distinguish co-expression within cells from shared expression across a tissue region. For cell types, co-localization curves could help distinguish proximity from occupancy of the same broad compartment, complementing existing methods [34–36], and extending these analyses to gene expression and pathways scores without requiring cell labels. Another potential Spectrast extension is to retain directional information when applying the Fourier transform, which can help describe highly structured or aligned tissue structures such as ducts, fiber tracts, or crypts, by direction and scale.

### 4.3 Limitations

While Spectrast makes multiscale analysis of high-resolution ST data possible, whole-transcriptome radii sweeps on large tissue sections are still very computationally expensive. Spectrast’s execution time and memory cost depends primarily on grid size, so large tissues measured at fine resolution will be the most expensive to analyze. Spectrast’s differential framework is also bounded by the number of tested biological replicates. Fewer samples limit the number of possible label reassignments when constructing the permutation-based null distribution, and therefore sets a lower bound on the minimum permutation p-value returned by the tool. We do not recommend reporting p-values for significance with less than six replicates per arm. Spectrast rasterizes ST data, which introduces approximation error. While this error is corrected in the current implementation using the cell’s original coordinates, Spectrast does not check the exact error for every cell and instead uses a proxy measure. The combination of rasterization and refinement improves estimates but will not always match the statistic value computed by the standard reference implementation. Accurate local measurements therefore depend on using a sufficiently small grid size. In most cases, Spectrast’s auto-tune feature automatically handles grid size determination. Other parameters apart from radius can affect spatial statistic values. Kernel shape, distance weighting, normalization, and spatial smoothing could also affect measured values, and the resulting scalogram. For example, inverse-distance weights give closer cells more influence compared to a binary weighting approach. We did not systematically evaluate these choices here, so it should not be assumed that results hold across all parameter settings. Finally, for large tissue sections, Spectrast’s reported statistical significance values can be uninformative. In some analyses, close to all gene-radius combinations will be significant, despite absolute spatial statistics being small. Effect sizes and scale profiles should be used alongside p-values to judge the biological relevance of Spectrast’s output.

### 4.4 AI disclosure

Generative AI was utilized for code generation, with human review and editing. Text was human written, with AI used for copy editing and grammar review.

## Data availability

All datasets analyzed in this study are publicly available. Quality-control filtering, cell and gene counts, and the full per-dataset processing steps are given in Supplementary Section S1.

- **Human ovarian cancer** (10x Visium HD), https://www.10xgenomics.com/datasets/visium-hd-three-prime-ovarian-cancer-discovery-fresh-frozen.
- **Coronal mouse brain** (10x Visium HD, CytAssist V4), https://www.10xgenomics.com/datasets/visium-hd-cytassist-gene-expression-libraries-of-mouse-brain-he-v4.
- **Sagittal mouse brain** (Vizgen MERFISH), Allen Institute Brain Cell Atlas, Zhuang-ABCA-3 section 3.010, accessed with the abc_atlas_access package (https://alleninstitute.github.io/abc_atlas_access/notebooks/zhuang_merfish_tutorial.html).
- **Mouse olfactory bulb** (Stereo-seq), https://github.com/JinmiaoChenLab/SEDR_analyses/tree/master/data.
- **Human breast cancer and human cervical cancer** (10x Atera whole-transcriptome preview), https://s3-us-west-2.amazonaws.com/10x.files/samples/atera/dev/WTA_Preview_FFPE_Breast_Cancer and https://s3-us-west-2.amazonaws.com/10x.files/samples/atera/dev/WTA_Preview_FFPE_Cervical_Cancer.
- **Mouse brain and human breast cancer** (10x Xenium Prime 5K), obtained from the 10x Genomics public dataset portal, https://www.10xgenomics.com/datasets.
- **Human reactive lymph node** (10x Xenium Prime 5K preview, FFPE), output bundle https://s3-us-west-2.amazonaws.com/10x.files/samples/xenium/3.0.0/Xenium_Prime_Human_Lymph_Node_Reactive_FFPE/Xenium_Prime_Human_Lymph_Node_Reactive_FFPE_outs.zip with the released cell-type labels https://cf.10xgenomics.com/samples/xenium/3.0.0/Xenium_Prime_Human_Lymph_Node_Reactive_FFPE/Xenium_Prime_Human_Lymph_Node_Reactive_FFPE_cell_types.csv.
- **Mouse brain *Trem2*** ^**R47H**^**/5xFAD** (Vizgen MERSCOPE MERFISH), Brain Image Library, /bil/lz/kjohnst2/b72faf9d87d7fc00 [43].
- **Human breast DCIS/IDC** (10x Xenium), Bioconductor **SubcellularSpatialData** via **ExperimentHub**, accession EH8567, sample IDC (https://www.bioconductor.org/packages/release/data/experiment/html/SubcellularSpatialData.html).
- **Mouse embryo E9.5–E16.5** (Stereo-seq, MOSTA) [44].

## Code availability

The toolkit is available at https://github.com/gingerii/spectrast.

